# Regions of Inflammation in mouse asthma correspond to regions of heme-free soluble guanylyl cyclase and can be tracked by marked expression of heme-oxygenase-1

**DOI:** 10.1101/2023.11.29.569245

**Authors:** Mamta Sumi, Rosemary Westcott, Eric Stuehr, Chaitali Ghosh, Dennis J. Stuehr, Arnab Ghosh

## Abstract

Asthma is characterized by airway remodeling and hyperreactivity. Our earlier studies determined that the Nitric Oxide (NO)-soluble Guanylyl Cyclase (sGC)-cGMP pathway plays a significant role in human lung bronchodilation. However this bronchodilation is dysfunctional in asthma due to high NO levels which cause sGC to become heme-free and desensitized to its natural activator, NO. In order to determine how asthma impacts the various lung segments/lobes we mapped the inflammatory regions of lungs to determine whether such regions coincided with molecular signatures of sGC dysfunction. We demonstrate using models of mouse asthma (OVA, CFA/HDM) that the inflammed segments of the mouse asthma lungs can be tracked by upregulated expression of HO1 and these regions in-turn overlap with regions of heme-free sGC as evidenced by a decreased sGC-α1β1 heterodimer and an increased response to heme-independent sGC activator, BAY 60-2770 relative to naïve uninflamed regions. We also find that NO generated from iNOS upregulation in the inflamed segments has a higher impact in developing heme-free sGC as increasing iNOS activity correlates linearly with elevated heme-independent sGC activation. This excess NO works by affecting the epithelial lung hemoglobin (Hb) to become heme-free in asthma thereby causing the Hb to lose its NO scavenging function and exposing the underlying smooth muscle sGC to excess NO, which in-turn becomes heme-free. Recognition of these specific lung segments enhance our understanding of the inflammed lungs in asthma with the ultimate aim to evaluate potential therapies and suggests that regional and not global inflammation impacts lung function in asthma.

## Introduction

Asthma is a chronic heterogeneous disease of the lower airways characterized by chronic inflammation and bronchial airway hyper-reactivity with variable airflow limitation (1, 2). The pathophysiology of asthma is complex and is estimated to impact 300 million people worldwide (2, 3). The specific bronchial airways that are impacted by asthma may vary in patients and between the types of asthma (3). While it is well understood that asthma presents differently in different airways, the disease is usually diagnosed using standard breath tests that measures the mean air flow in all the airways (4). This type of diagnostic approach overlooks sub regions of the disease where localized inflammation of the airways maybe more prevalent and targeted therapeutics may be essential to achieve localized dilation. Inflammation in the lung is the body’s natural response to injury and acts to remove harmful stimuli such as pathogens, irritants or damaged cells and initiate the healing process. Both acute and chronic inflammation are seen in different respiratory diseases such as; acute respiratory distress syndrome, chronic obstructive pulmonary disease (COPD) and asthma. Inflammation in asthma can be patterned differently depending upon the onset of disease progression. Chronic asthma and late asthmatic responses (LAR) are associated with local inflammation which might be expected to produce airflow obstruction in small airways and to increase nonspecific airway reactivity. In contrast, early asthmatic responses (EAR) are primarily bronchospastic and probably involve more central airways (2, 5). The molecular underpinnings associated with such inflammatory changes is incompletely understood and studying these aspects would allow us to better understand the inherent drivers of these processes. Using translational models of mouse asthma (6) it is possible to study the distribution of inflammation in all segments (7) of the asthma lungs with the ultimate aim to improve patient treatments and to evaluate potential therapies.

Although airway inflammation plays a role in severe asthma, inherent dysfunction in the airway epithelial and/or smooth muscle cells, which are pivotal in regulating bronchomotor tone, may also contribute to the asthma diathesis (3, 8–11). Human airway smooth muscle cells (HASMC) play a central role in bronchomotor function and in asthma their dysregulation promotes the airway narrowing, obstruction and hypersensitivity that characterizes the disease (9, 12). The NO-sGC-cGMP pathway is the primary driver of vascular smooth muscle relaxation (13–15), and although its role in HASMC relaxation has long been suspected (16), it was only recently established that activating this pathway is as effective in evoking bronchodilation in human small airways as is activating the β_2_AR-sAC-cAMP pathway (17, 18). However sGC becomes dysfunctional in allergic asthma due it becoming heme-free on account of excess NO generated from iNOS upregulation during lung inflammation, which makes the epithelia lung hemoglobin (Hb) heme-free (19), thereby making it unable to scavenge this excess NO and in-turn exposing the underlying smooth muscle sGC to excess NO, triggering its dysfunction (19, 20). In order to understand whether such a sGC dysfunction is associated with inflammation which is either localized or globally distributed in an asthma lung we resorted to connect the inflammatory regions to sGC dysfunction using mouse asthma (Ovalbumin [OVA] or Complete freund’s adjuvant/House dust mite [CFA/HDM]) lungs which underwent lobar segmentation into 14 lung segments (7). Overall objective of the present study was to monitor the dynamics of inflammatory changes taking place in an asthmatic mouse lung and determine i) the extent of localized inflammation, ii) whether localized inflammatory regions can be correlated with sGC dysfunction by determining the heme status of the sGC in such areas of inflammation, iii) whether such inflammatory areas can be tracked with molecular indicators of inflammation and iii) whether such regions in-turn have increased iNOS activity since high NO levels have been shown to cause sGC dysfunction in asthma. To decipher whether all these parameters overlap on a fulcrum point of high inflammation we first subjected naïve, OVA or CFA/HDM asthma mice lungs to undergo lobar segmentation and then use the generated lung tissue supernatants from the individual 14 sections of each mouse lung to perform biochemical assays comprising of western blots, immunoprecipitation assays, enzyme activity assays, heme-stains/heme estimations and tissue immunostaining.

## Materials and Methods

### Reagents

All chemicals were purchased from Sigma (St. Louis, MO) and Fischer chemicals (New Jersey). TNFα, Human interferon gamma (IFN-γ), interleukin 1β (IL-1β), Phosphodiesterase inhibitor 3-isobutyl-1-methylxanthine (IBMX), *o*-Dianisidine for heme-staining, sGC stimulator BAY 41-2272 (BAY-41, heme-dependent) and activator BAY 60-2270 (BAY-60, heme-independent) were purchased from Sigma. Heme estimation kit was purchased from Abnova. Reagents such as L-Arginine, Sodium pyruvate, Lactate dehydrogenase (LDH), NADPH, flavins (FAD, FMN), Tetrahydrobiopterin (H_4_B) needed for NOS activity assays were also obtained from Sigma. 16-HBE cells were purchased from American Type Culture Collection (ATCC; Manassas, VA, USA). cGMP ELISA assay kits were obtained from Cell Signaling Technology (Danvers, MA, USA). Protein G-sepharose beads were purchased from Sigma and molecular mass markers were purchased from Bio-Rad (Hercules, CA, USA).

### Antibodies

Antibodies were purchased from different sources. Supplemental Table S1 describes various types of antibodies used and its source.

### Growth/induction of cells, Western blots and immunoprecipitations (IPs), heme estimations and heme-stains

All cells were grown and harvested as previously described (20). 16-HBE bronchial epithelial cells were cultured to confluency (50-60%) and then induced with TNF α, (20 ng/ul), IFN-γ (10 ng/ul) and IL-1β (10 ng/ul) from 0-48h. Cell supernatants were prepared at different time points at 0, 12, 24, 36 and 48h to analyze for induced protein expression by western blots and protein-protein interactions by immunoprecipitation assays (IPs). Culture media was also collected in parallel to assay for accumulated nitric oxide (NO). Western blots were performed using standard protocols as previously mentioned. For westerns involving multiple samples 50-80ug of the lung supernatants from naïve, OVA or CFA/HDM mice were run separately on large SDS-PAGE (8%), transferred to the same PVDF membrane, probed with a specific antibody and developed at the same time. β-actin was used as a loading control. Multiple protein detection was achieved by stripping the membranes and re-probing with specific antibodies. For immunoprecipitations (IPs), 400-600 μg of the total cell supernatant was precleared with 20 µl of protein G-sepharose beads (Amersham) for 1 h at 4° C, beads were pelleted, and the supernatants incubated overnight at 4 °C with 3 μg of the indicated antibody. Protein G-sepharose beads (20 µL) were then added and incubated for 1 h at 4 °C. The beads were micro-centrifuged (6000 rpm), washed three times with wash buffer (50 mM HEPES pH 7.6, 100 mM NaCl, 1 mM EDTA and 0.5% NP-40) and then boiled with SDS-buffer and centrifuged. The supernatants were then loaded on SDS-PAGE gels and western blotted with specific antibodies. Band intensities on westerns were quantified using Image J quantification software (NIH). Heme-estimations on mouse lung tissue supernatants were done using the heme-estimation assay kit, while heme-stains on such supernatants were performed as earlier described (21).

### Mouse OVA and CFA/HDM Models of Asthma

The mouse OVA model are characterized by eosinophilic asthma, develops airway hyperreactivity (AHR), goblet cell metaplasia and is reactive to steroid treatment (20, 22–24). Allergic airway disease was performed following procedures as described earlier (25, 26). 12-weeks old female C57BL/6 mice from Jackson Laboratory (Bar Harbor, Me) were used for OVA allergen sensitization and and challenge. All mice experiments were approved by the Cleveland Clinic Institutional Animal Care and Use Committee. Mice were sensitized by intraperitoneal injection with OVA (Sigma Chemicals) [10 µg, adsorbed in Al [(OH)_3_] on days 0 and 7 and challenged with aerosolized OVA (1% w/v in sterile PBS) on days 14-19, which included a series of daily inhalations which lasted 40 min/day, where the mice were placed in a chamber kept saturated with nebulized OVA solution (1% w/v in sterile PBS). Animals were processed on day 20. The animals were anesthetized by means of intraperitoneal injection with pentobarbital and bronchoalveolar lavage (BAL) fluid collection for cell counting or tissue collection were performed 24 hours after the last OVA challenge. On the collected BAL, differential counts for eosinophils, lymphocytes, neutrophils, or alveolar macrophages were performed. All counts were performed by a single observer blinded to study groups. The lungs were then dissected and segmented into 14 sub-lobar segments (number of segments per lobe: left 4, accessory 2, inferior 4, intermediate 2 and superior 2) for biochemical analyses.

The CFA/HDM model is a neutrophilic asthma model, develops airway hyperreactivity (AHR), goblet cell metaplasia and is steroid-resistant (27–31). The CFA/HDM acute asthma model was done as described previously (29). Here 12-weeks-old female WT C57BL/6 mice (from Jackson Laboratory) were sensitized subcutaneously on day 0 with HDM (*D.Farinae*) extract (100 μg per mouse) (32) emulsified in CFA (Greer labs) and subsequently challenged intranasally after 14 days with HDM (100 μg in 50 μl saline per mouse). The endotoxin level in purchased HDM was in the range of 2.9 x 10^3^ to 9.7 x 10^3^ EU/mg as provided by Greer labs. BAL for cell counting and lung tissue collection were performed 24 hours after the last HDM challenge similar to that described for the OVA model. Parallel studies on Airway Hyper-responsiveness (AHR) and Lung mechanics were measured using the FlexiVent ventilator (FlexiVent, Scireq) 24 h after the last OVA/HDM challenge in response to increasing doses of inhaled methacholine (Mch) (i.e from 0, 25, 50 and 100mg/ml doses of Mch) that was used as a bronchoconstrictor and quantifications were done as previously described (33).

### Immunohistological staining of mice lung tissues

Formalin-fixed paraffin embedded (FFPE) lung tissue sections from naïve or OVA mice were immunostained following standard protocols. Tissue sections were de-paraffinized and rehydrated by immersing the slides through three different solvent washes. Three washes of xylene for 5 minutes each, ethanol (100%/95%/70%/50%, for 10 min each) and water (10 mins each) were done before antigen retrieval by tris EDTA (pH 9). PBS blocking solution (including 1% Triton X-100 and 10% goat serum) was used to block sections and as a diluent for primary and secondary antibodies. Primary antibodies were rabbit polyclonal antibody against heme-oxygenase-1 (Cell Signaling Tech., 1:20 dilution) and mouse monoclonal antibody against iNOS (Invitrogen, 1:20 dilution) were used. Secondary antibodies used were donkey anti-mouse (Alexa Fluor 488, Jackson Immunoresearch, 715-545-150, 1:200 dilution) and donkey anti-rabbit (Texas red, Jackson Immunoresearch, 711-076-152, 1:200 dilution). Stained sections were then mounted, and imaged on a Zeiss Axio vertA1 microscope or confocal microscopy was performed. *DAB Staining:* Immunohistochemistry staining was performed using the Discovery ULTRA automated stainer from Roche Diagnostics (Indianapolis, IN). In brief, antigen retrieval was performed using a tris/borate/EDTA buffer (Discovery CC1, 06414575001; Roche), pH 8.0 to 8.5. Time, temperatures, and dilutions are listed below. The antibodies were visualized using the OmniMap anti-Rabbit HRP (05269679001; Roche) in conjunction with the ChromoMap DAB detection kit (05266645001; Roche). Lastly, the slides were counterstained with hematoxylin and bluing. *H& E Staining* was performed on mouse lung sections as described previously (34).

### Preparation of lung tissue supernatants

Mice Lung tissue segments from naïve, OVA or CFA/HDM lungs were processed to generate tissue supernatants. These tissue samples were washed extensively (6 times) with RBC lysis buffer (Cell Signaling Technology) until the soret absorption peak on the UV-visible spectra was almost negligible and the washes were colorless. This ensured that the tissues were free from residual blood in its capillaries. Tissue supernatants were then prepared on the washed samples.

### Olis Clarity spectroscopy

The analysis of Hb heme in the supernatants of CFA/HDM lung segments was done by using the Olis Clarity spectrophotometer (21), which can record the heme spectra of suspensions or solutions in particulate form. Here the absorption spectra was collected with an integrating sphere detector. Segments of mice lungs from naïve and CFA/HDM mice (25, 26, 33) were first washed with 1X PBS to remove residual blood sticking to the capillaries of the lung tissues and then were washed extensively (6 times) with RBC lysis buffer. IPs were performed on 400 ug of total protein with Hbβ antibody, beads washed with wash buffer and the bead bound Hbβ spectra from lung tissue supernatants were recorded on the Olis Clarity.

### cGMP enzyme-linked immunosorbent assay

Lung tissue supernatants from naïve, OVA or CFA/HDM mice were assayed for sGC enzymatic activity (20, 35). Reactions containing aliquots of the tissue supernatants supplemented with 250 μM GTP, 10 μM of sGC stimulator BAY-41 or activator BAY-60, 250µM IBMX were incubated for 20 min at 37 °C. Reactions were quenched by addition of 10 mM Na_2_CO_3_ and Zn (CH_3_CO_3_)_2_. The generated cGMP was then estimated by the cGMP ELISA assay kit (Cell Signaling Technology).

### In vitro NOS reconstitution and Nitrite estimation in the culture media

iNOS activity in mouse lung supernatants was determined by measuring production of nitrite alone or nitrite plus nitrate (stable oxidation products of NO that accumulate quantitatively) in 30-min incubations run at 37°C. Aliquots of the mice lung supernatants were transferred to microwells containing 40 mM Tris buffer (pH 7.8) supplemented with 3 mM DTT, 2 mM l-arginine, 1 mM NADPH, protease inhibitors, and a 4 mM concentration each of FAD, FMN, and H4biopterin, to give a final volume of 0.1 ml. Reactions were terminated by enzymatic depletion of the remaining NADPH (36). In separate experiments the cell culture media from induced 16-HBE cells expressing iNOS were taken out at specific time points to determine the accumulated nitrite. Nitrite was measured using ozone-based chemiluminescence with the triiodide method and using the Sievers NO analyzer (GE Analytical Instruments, Boulder, CO, USA) as described earlier (37, 38).

## Results

### Expression of specific proteins and lowered tissue heme levels in mouse asthma (OVA) lungs

In a previous study (39) we found that expression of sGCβ1, key redox proteins eg. catalase, Trx1 etc. were downregulated while heme catabolic enzymes HO1, HO2 were upregulated in human airway smooth muscle cells from severe asthma. We did expression profiling of these proteins in lung tissue supernatants from mouse asthma (OVA) and determined a like pattern whereby sGCβ1, catalase, trx1 were downregulated, while heme-oxygenases were upregulated relative to naïve controls (Figs. 1A, B), suggesting that similar regulatory mechanisms of gene expression may operate in mouse asthma. Our earlier study (20) found that lung bronchodilator protein sGC, was dysfunctional in asthma as it became heme-free and cannot be activated by its natural activator NO leading to obstructed bronchodilation. This led us to investigate the overall heme levels of hemeproteins in mouse lung supernatants from asthma relative to naïve controls. As depicted in Fig. 1C we found that total heme-stains were lowered in the mouse OVA lungs and the mean heme content was almost two fold lowered relative to naïve lungs as quantified by heme-estimation (Fig. 1D). Together our data suggests that asthma also negatively impacts the heme levels on hemeproteins other than sGC and this maybe attributed to upregulated levels of heme oxygenases.

**Figure 1.**
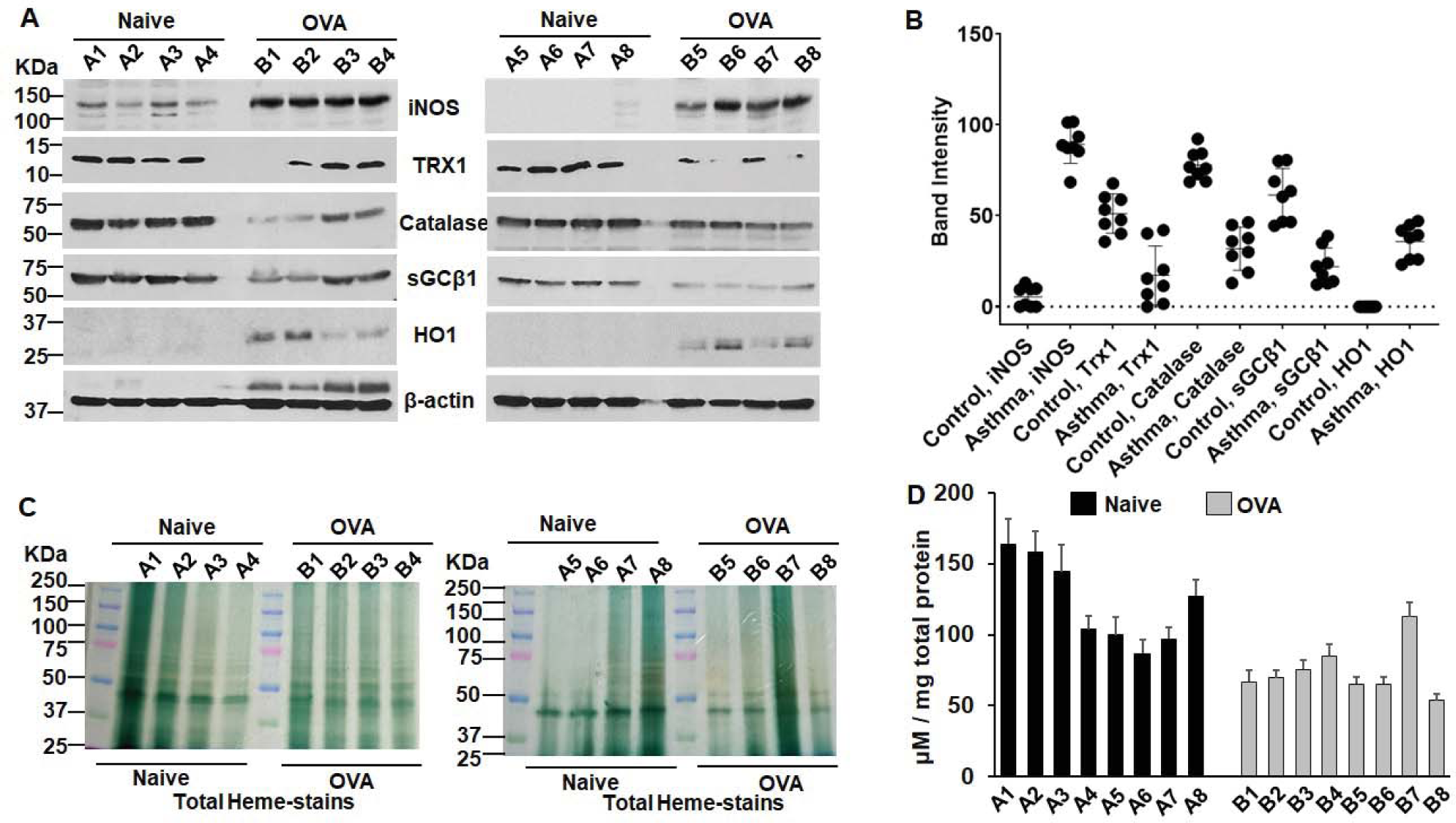
Lung expression of signal, redox and heme catabolic proteins/enzymes in a mouse asthma model. Mouse lung supernatants from control naïve and asthma model, OVA were run on 8 or 15% SDS-PAGE and western blotted with specific antibodies or heme-stained. Heme-estimations on total lung supernatants were also performed. (A) Representative expression of proteins by westerns as depicted. (B) Densitometries of protein expression shown in A. Values depicted are mean densitometry values, -/+ SEM from n=8 mice lungs. (C) Heme-stains of total protein generated from mouse lung supernatants of naïve and OVA mice. (D) Total heme concentrations in the mouse lung supernatants calculated using a heme-estimation assay kit. Data are mean (n= 3 repeats per condition) ± SD. Wherever applicable molecular weight markers (KDa) are depicted at the left of gel bands throughout the figure legends.

### Select expression of HO1 in inflammatory regions of mouse asthma (OVA) lungs

Figure 2A. outlines the segmentation of the various lung lobes of the mouse and the overall experimental plan. We first needed to determine that whether inflammation was uniform in all the 14 lung segments of the asthma mice lungs. We used the OVA mice lung segments to determine the extent of inflammation in the asthma mice lungs and we probed for the expression of a inflammatory protein eg. HO1 in the various lung segments (Fig. 2) and compared it to naïve control lungs. As depicted by western expression and corresponding heat maps/densitometries in fig. 2 or supplemental fig. S1, we found variable expression of HO1 in the different asthma mouse (B1 to B5) and these were elevated in specific lung segments which maybe more inflammed relative to less inflammed or non-inflammed (naïve mice lung segments) regions. This may result from non-uniform inflammation occurring in various lung segments of asthma mice causing variable readout of HO1 expression. From figs. 2B, C and supplemental fig. S1, we determine that HO1 was specifically elevated in S1, L1, L3, A1 for B1; S1, S2, IF2, L1, A1 for B2; S1, S2, Int1, IF2, L1, L3, A1 for B3; Int1, IF2, L1, A1 for B4; IF2, L1, L3, A1 for B5. Among all these asthma mice L1, L3 and A1 regions were the most common to show elevated expression of HO1, S1/S2 and IF2 were the next most common while Int1 was also found elevated in some mice (Fig. 2B & C). However the other heme-oxygenase HO2 which is constitutively expressed (39, 40) did not display such specificity of expression in the inflammatory regions like HO1 and only a sparsely elevated expression prevailed in the A1 region (Fig. S1). Together these data imply that specific inflammatory regions of the asthma (OVA) mice lungs display enhanced expression of HO1.

**Figure 2.**
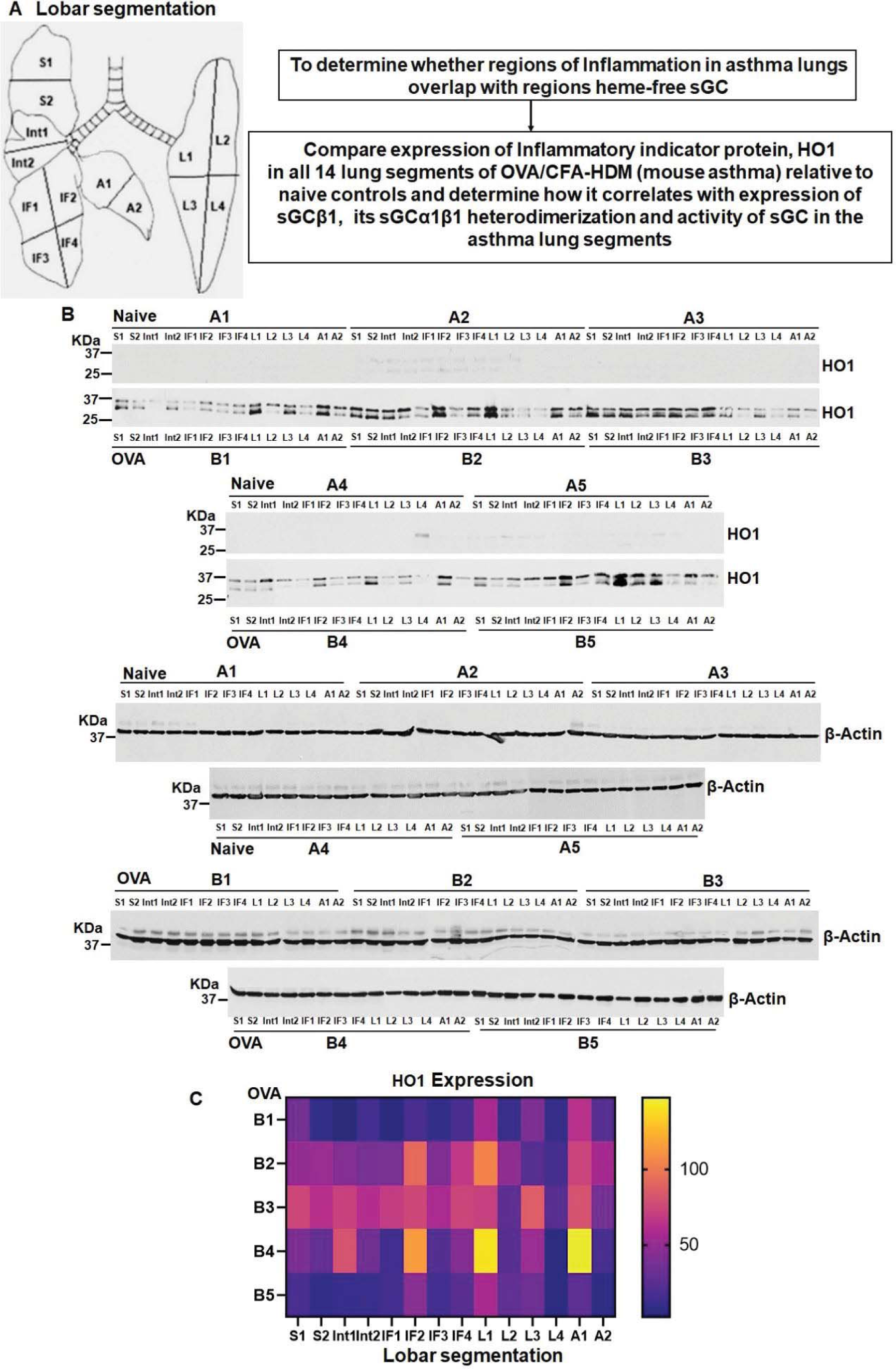
Expression of HO1 in the lung supernatants from control naïve and asthma (OVA) mice. Mouse lung supernatants generated from all the 14 segments of a mouse lung were run on SDS-PAGE and western blotted with specific antibodies as depicted. (A) Lobar segmentation [adapted from Asosingh et al. (7)] and Experimental design. (B) Representative expression of HO1 in naïve (A1-A5) and OVA (B1-B5) mice lung segments with β-actin used as a loading control. (C) Heat map of HO1 expression (as depicted in panel B) derived from corresponding densitometries as depicted in figure S1.

### Regions of inflammation can be tracked by elevated expression of HO1 which overlaps with regions of heme-free sGC in mouse asthma lungs

To determine the expression of sGCβ1 we did westerns in the lung segments of naïve and OVA mice and as depicted from the blots or its corresponding heat maps/densitometries in Fig. 3A, B or supplemental fig. S2, there was lowered expression of sGCβ1 in the asthma mouse relative to naïve controls. Doing IPs to determine the status of the lung sGC heterodimer, we choose four segments from each mouse OVA lung which were upregulated in HO1 and one that was least. The IPs revealed that sGCα1β1 heterodimer was very low (Fig. 3C) in the highly expressed HO1 segments of the mouse OVA lungs and plotting the specific HO1 expression densitometries (from Fig. 2 and S1) versus the sGC heterodimer densitometries revealed an inverse correlation for each of the five asthma lungs (Fig. 3C, D). Doing sGC activation assays by activating the two populations of sGC (heme-free and heme-containing forms) with heme-independent (BAY-60) and heme-dependent (BAY-41) sGC activators we found that most inflammed regions which were marked by elevated expression of HO1 were heme-free as depicted by enhanced activation by BAY-60 (Fig. 3E). Plotting the HO1 densitometries versus the ratio of BAY-60/BAY-41 for all 14 lung segments of OVA mice revealed a direct correlation as increased HO1 expression caused increased heme-free sGC (Fig. 3F). In order to further establish how universal these findings were in other models of mouse asthma, we used mouse lung segment tissue from a severe asthma model eg. CFA/HDM (Figs. 4 and S5), which is different from the eosinophilic OVA model (20) as it is predominantly neutrophilic (Fig. S6). Probing for HO1 expression in the mouse lung segments we found a similar pattern of elevated HO1 expression in the asthma mouse relative to the naïves, and a variable expression of HO1 in the different lung segments was also found but with some differences in the segments showing upregulated expression relative to the OVA lungs (Figs. 4A, B and S3). Here we determined that segments S2, IF3, IF4 and A1/A2 had higher HO1 expression relative to other regions which was non-uniformly distributed over the five asthma mice (B1-B5, Figs. 4A, B and S3). We also determined a similar lowering of sGCβ1 expression in these asthma mice relative to the naives like we earlier found for the OVA model (Figs. 4A, B and S3). Assaying for the status of sGCα1β1 heterodimer we found that the chosen three out of five regions which displayed elevated HO1 expression had a poor sGC heterodimer while the other two regions which were low on HO1 expression showed a relatively better heterodimer and plotting the HO1 densitometries versus the sGC heterodimer displayed an inverse correlation (Figs. 5A-C). sGC activation assays also showed a predominant response to heme-independent sGC activator BAY-60 in regions showing elevated HO1 expression (Fig. 5D), with a linear correlation existing between HO1 expression and BAY-60/BAY-41 ratios (Fig. 5E). Together our data from both asthma models depicts that inflammatory regions of asthma lungs which are characterized by marked expression of HO1 have a poor sGC heterodimer on account of sGCβ1 being heme-free.

**Figure 3.**
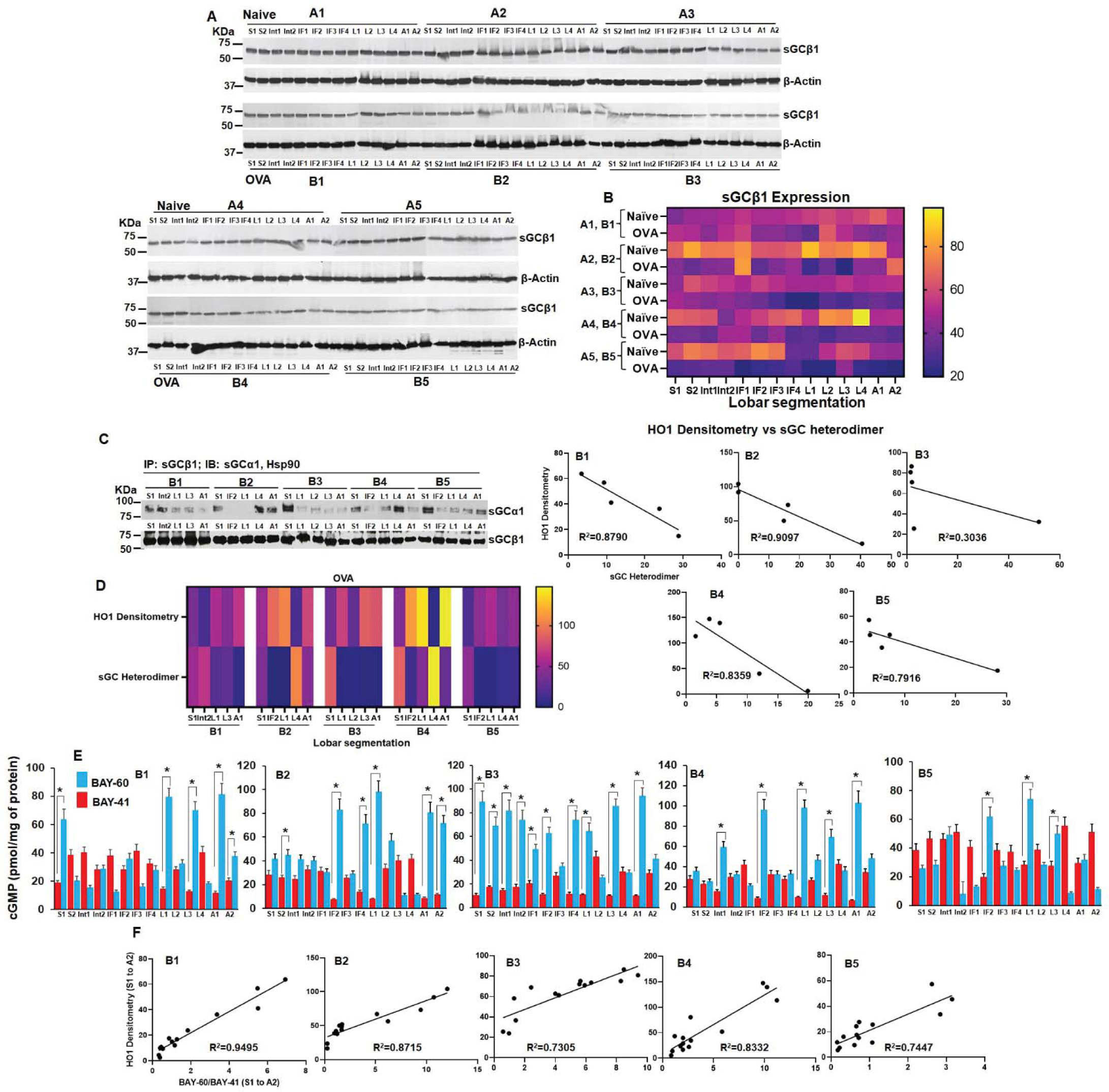
Points of upregulated HO1 expression develop more heme-free sGC which is characterized by a poor sGC heterodimer and is BAY-60 responsive. Mouse lung supernatants from naïve and OVA mice lung segments underwent western blotting for sGCβ1 expression, immunoprecipitation assays (IPs) and sGC activation assays with BAY-41 and BAY-60. (A) Representative expression of sGCβ1 by westerns. (B) Heat map of sGCβ1 expression derived from its corresponding densitometries in naïve and OVA mice lung segments as depicted in figure S2. (C) IPs depicting sGC heterodimer and its inversely correlated densitometries of HO1 expression and sGC heterodimer in the mouse OVA lung segments (n=5 mice). (D) Heat map depicting inverse correlation of HO1 expression and sGC heterodimer from corresponding densitometries in OVA lung segments. (E) sGC activation assays with BAY-41 or BAY-60, with ELISA as a readout to estimate the generated cGMP. (F) Densitometries of HO1 expression directly correlates with sGC activation with BAY-60. Lines of best fit are shown along with the correlation coefficient. Data are mean (n= 3 repeats per condition) ± SD. *p < 0.05, by one-way ANOVA.

**Figure 4.**
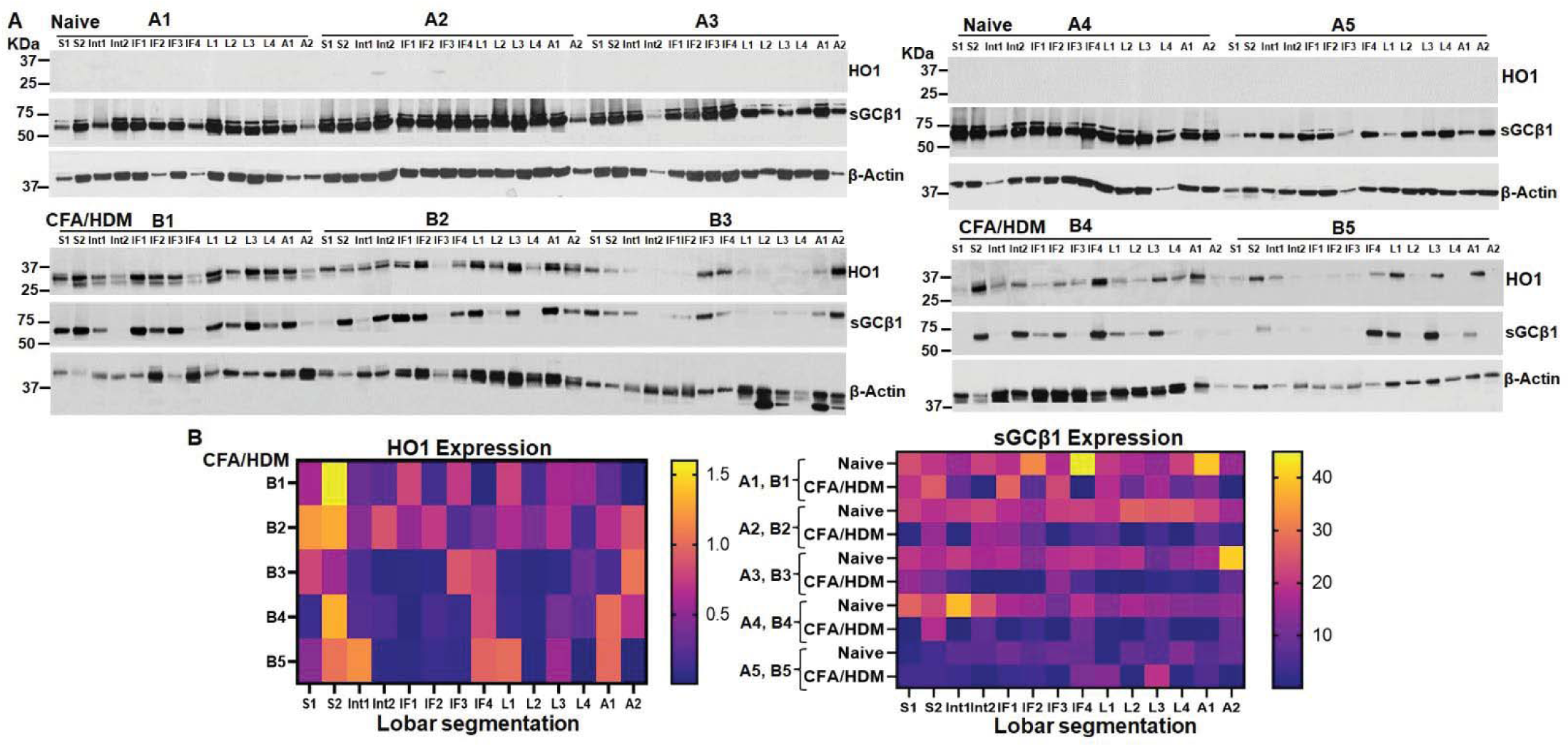
Expression of HO1 and sGCβ1 in the lung supernatants from control naïve and asthma (CFA/HDM) mice. Mouse lung supernatants generated from all the 14 segments of a mouse lung were run on SDS-PAGE and western blotted with specific antibodies as depicted. (A) Representative expression of HO1 and sGCβ1 in naïve (A1-A5) and CFA/HDM (B1-B5) mice lung segments with β-actin used as a loading control. (B) Heat maps of HO1 and sGCβ1 expression (panel A) in naïve and CFA/HDM mice lung segments derived from corresponding densitometries as depicted in figure S3.

**Figure 5.**
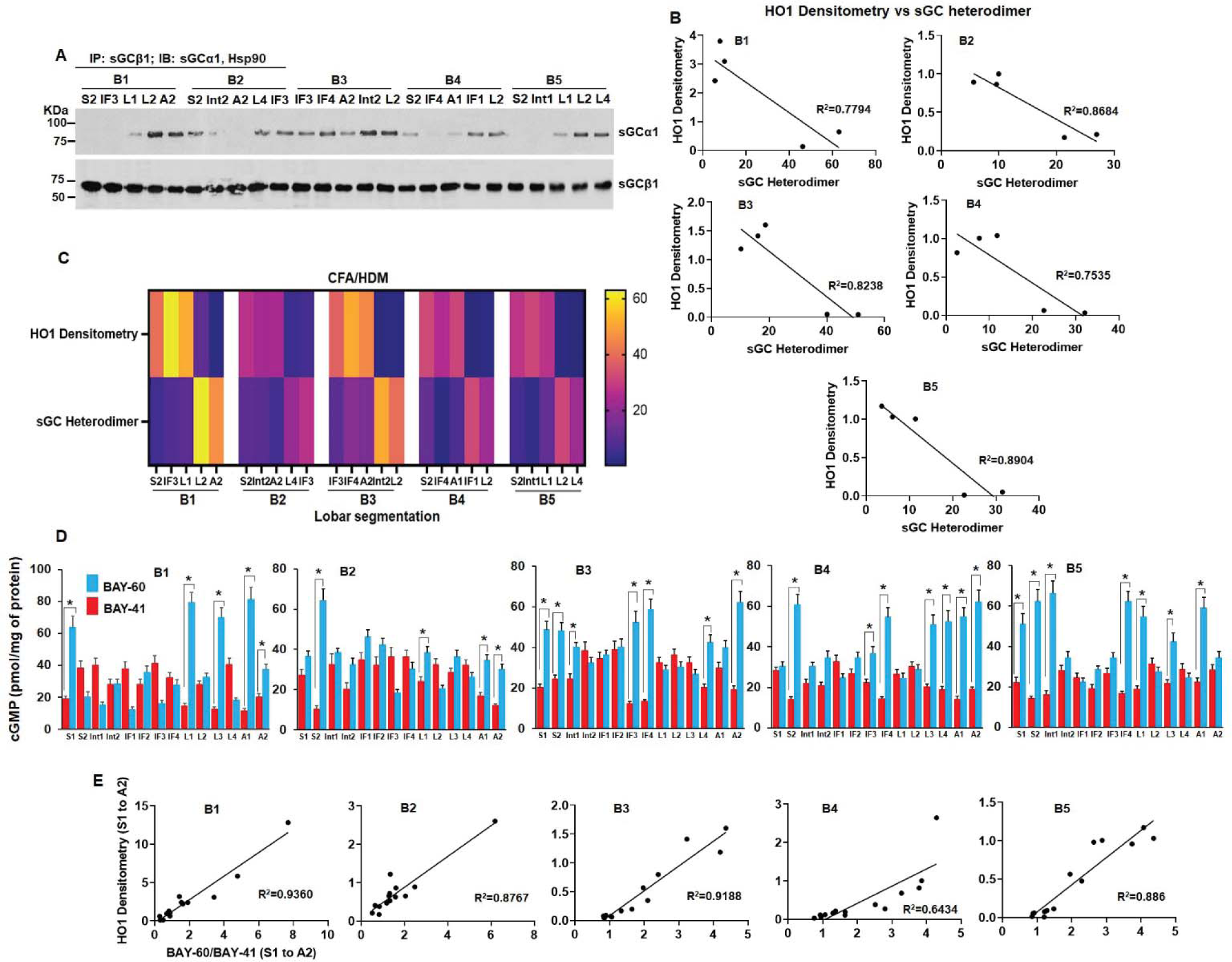
Regions of upregulated HO1 expression in CFA/HDM lung segments corresponds to heme-free sGC which is characterized by a poor sGC heterodimer and is BAY-60 responsive. Mouse lung supernatants from naïve or CFA/HDM mice lung segments underwent immunoprecipitation (IPs) and sGC activation assays with BAY-41 and BAY-60. (A) IPs depicting sGC heterodimer. (B) Inversely correlated densitometries of HO1 expression and sGC heterodimer in the mouse CFA/HDM lung segments (n=5 mice). (C) Heat map depicting inverse correlation of HO1 expression and sGC heterodimer from corresponding densitometries in CFA/HDM lung segments. (D) sGC activation assays with BAY-41 or BAY-60, with ELISA as a readout to estimate the generated cGMP. (E) Densitometries of HO1 expression directly correlates with sGC activation with BAY-60. Data are mean (n= 3 repeats per condition) ± SD. *p < 0.05, by one-way ANOVA.

### iNOS activity correlates linearly with increase in heme-free sGC and upregulated HO1 expression

In order to attribute the reasons for developing heme-free sGC we determined the expression of iNOS in the various lung segments. As depicted in Fig. 6A, there was iNOS expression in the lung segments of OVA mice and iNOS reconstitution assay performed on these lung segment supernatants (Fig. 6B) revealed a variable pattern of NO generation which was more in specific regions that were marked with a higher HO1 expression. Plotting the iNOS activity against the corresponding HO1 densitometry in all lung segments of the five OVA mice showed a direct correlation (Fig. S4) suggesting that higher points of inflammation can be marked both by elevated expression of HO1 and a higher NO generating activity from the reconstituted iNOS. As high NO is one of the putative reasons for sGC becoming heme-free in asthma (20), plotting iNOS activity versus heme-independent sGC activation with BAY-60 gave a linear correlation where higher NO generating activity of iNOS coincided with greater generation of heme-free sGC (Fig. 6C). Cumulative heat maps of HO1 expression, iNOS activity and sGC activation by BAY-60 also depicted these correlations (Fig. 6D). Together these data suggest that points of inflammation in mouse asthma lungs have high NO generation from induced iNOS which can be tracked by elevated expression of HO1 and these regions in-turn have greater dysfunctional sGC that is heme-free.

**Figure 6.**
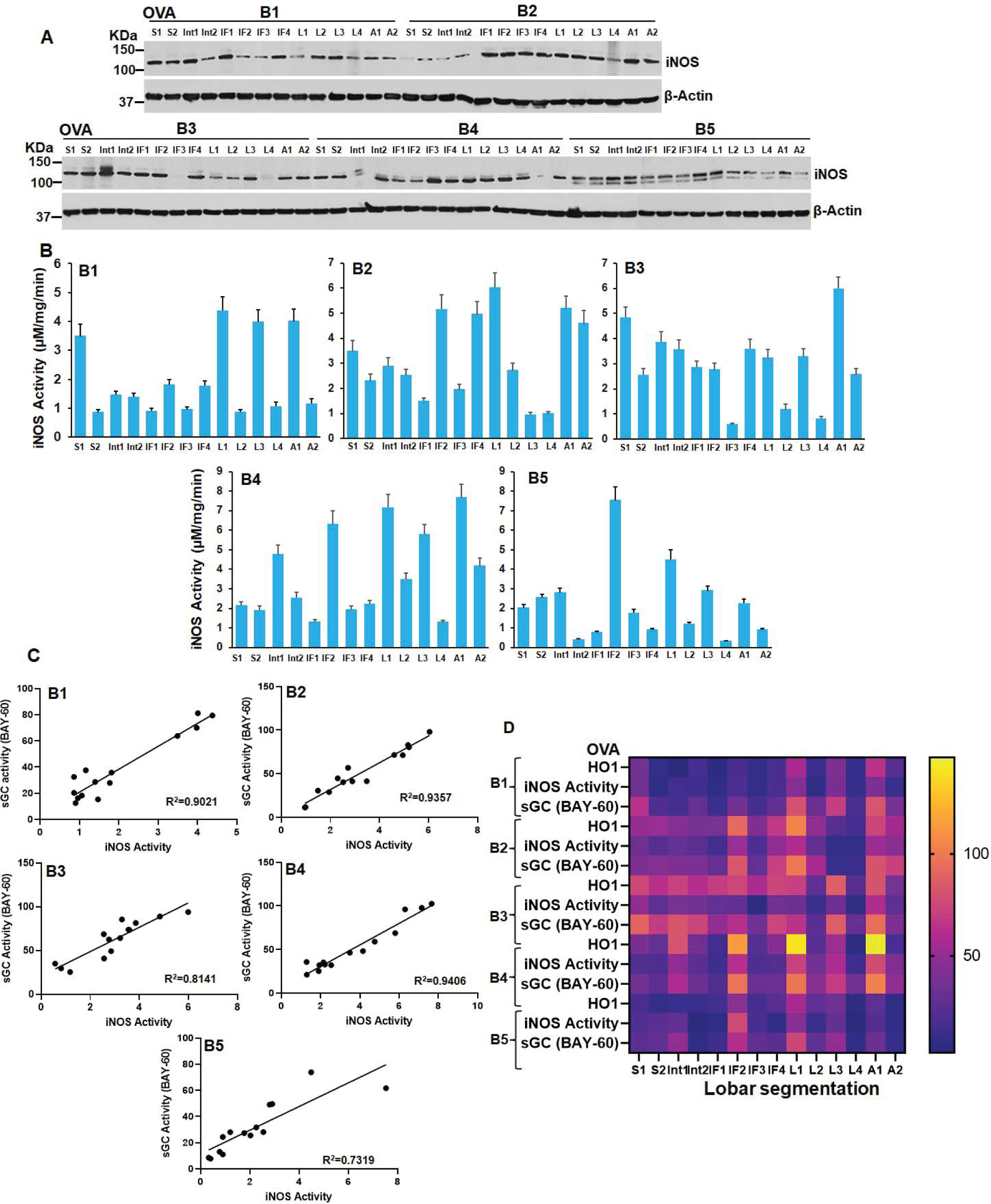
iNOS activity correlates linearly with increase in heme-free sGC and upregulated HO1 expression. Mouse lung supernatants from OVA mice lung segments underwent western blotting for iNOS expression and iNOS reconstitution assays. (A) Representative expression of iNOS by westerns in OVA mice lung segments with β-actin used as a loading control. (B) iNOS reconstitution assays using mouse lung supernatants from OVA mice. (C) Correlations between heme-independent sGC activation (BAY-60) and iNOS activity in OVA mice lung segments. Data are mean (n= 3 repeats per condition) ± SD. (D) Heat map depicting HO1 expression, iNOS activity and sGC activation (BAY-60) in OVA mice lungs.

### iNOS and HO1 interact or co-localize in bronchial epithelial cells and in mouse asthma

Since HO1 is known to modulate iNOS activity (41, 42) we tested the ability of these two proteins to interact in cytokine stimulated bronchial cells or in mouse asthma. As depicted in Fig. 7A, B, HO1 and iNOS showed parabolic pattern of interaction in cytokine stimulated 16-HBE cells, where time points of maximum interaction (24-36h, Fig. 7B) correlated with the surge in NO production from iNOS (Fig. 7C). This interaction was also significant in inflammed regions of OVA lungs which was marked by HO1 upregulation (Fig. 7D), and iNOS bound HO1 correlated inversely to the sGCα1β1 heterodimer (determined from Fig. 3C) in all five mouse OVA lung sections (Fig. 7D). Immunohistological staining in form of H & E staining revealed that there was narrowing of the airways (43) on account of inflammation in OVA lungs relative to the naïves (Fig. S7). While DAB staining and fluorescence imaging of OVA sections showed that iNOS and HO1 are induced in asthma lungs (Fig. 7E, F and G). Both HO1 and iNOS co-localize (Fig. 7H) which suggests that this interaction is vital to modulate iNOS activity as it can regulate the iNOS NO synthesis (Fig. 7C) which in-turn cascades into generation of heme-free sGC in the underlying smooth muscle cells. Since we also found that HO1 failed to interact with sGCβ1 (data not shown) the generation of heme-free sGC may be largely caused due to the impact of high NO generation in mouse asthma.

**Figure 7.**
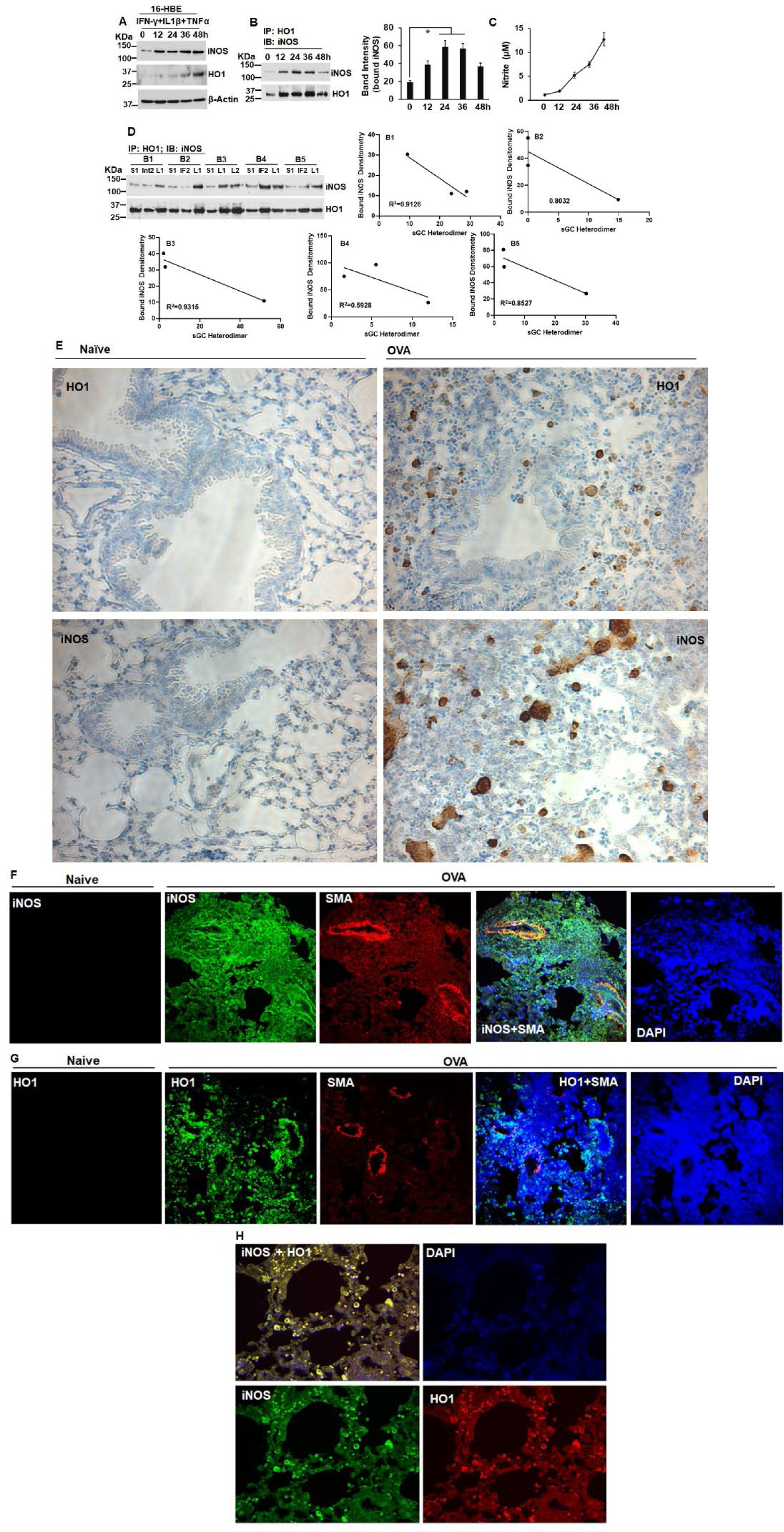
iNOS and HO1 interact or co-localize in bronchial epithelial cells and in mouse asthma. iNOS in the bronchial epithelial cells (16-HBE) was induced by cytokines for NO generation and to determine its interaction with co-induced HO1. In parallel experiments supernatants from OVA mice lung segments were tested for iNOS-HO1 interaction with IPs and immunohistological staining to study the expression or co-localization of these two proteins. (A) Western blots of protein expression in the 16-HBE cells as indicated. (B) IPs and its corresponding densitometries depicting iNOS-HO1 interaction between 0-48 h of cytokine induction of 16-HBE cells. (C) Accumulated NO in the culture media estimated as nitrite. (D) IPs depicting iNOS-HO1 interaction in OVA mice lung segments and its corresponding densitometries correlated with sGC heterodimer determined in the same lung segments of OVA mice as depicted in figure 4B (n=5, OVA mice). (E) DAB staining depicting positive expression of HO1 and iNOS in sections from OVA lungs relative to naïve lungs. (F-H) Images captured on a confocal microscope at 20x (F,G) and 40x (H) depicting iNOS or HO1 localization relative to smooth muscle actin (SMA) and iNOS-HO1 co-localization in mouse OVA lung sections. Values depicted are mean (n= 3 repeats per condition) ± SD. *p < 0.05, by one-way ANOVA.

### Heme status of Hb in asthma mouse lung segments

Our current study suggests that asthma negatively impacts the heme levels of hemeproteins in the asthma lungs (OVA) relative to naïve lungs with overall lowered tissue heme levels in asthmatic (OVA) lungs (Fig. 1). In this context our earlier study (19) described the role of Hb present in the lung epithelium and we found that it protects the underlying smooth muscle sGC by scavenging the NO. In both OVA and HDM asthma mice, excess NO generated largely from iNOS (20) can negatively impact the heme of lung Hb. In OVA lungs we found that this lung Hb becomes heme-free, thereby losing its NO scavenging function (19). We therefore determined the heme status of lung Hb in the CFA/HDM lung segments which predominantly gave a poor sGC heterodimer and displayed an elevated expression of HO1 (Fig. 5). As depicted in fig. 8A, we found that Hbβ immunoprecipitates from CHA/HDM lung segments retained a greater amount of hsp90 relative to naives, which suggests that the Hb heme was relatively heme-free. Measuring the Olis Clarity spectra on the bead bound Hbβ immunoprecipitates confirmed that Hbβ was heme-free relative to the naïve segments (Fig. 8B). Together these data suggests that lung Hb in asthma lungs becomes heme-free and is unable to protect the downstream sGC from adverse effects of excess NO (19). *Working model:* Based on our results from this study we construct a model which incorporates the molecular signatures of inflammation including HO1 upregulation, heme status of lung Hb and sGC dysfunction and shows that these converge on a common axes (Fig. 9) which are distributed in specific lung segments of individual asthma mice. Here high NO generation resulting from iNOS upregulation leads to heme-free sGC generation and these points can be tracked by upregulated expression of HO1.

**Figure 8.**
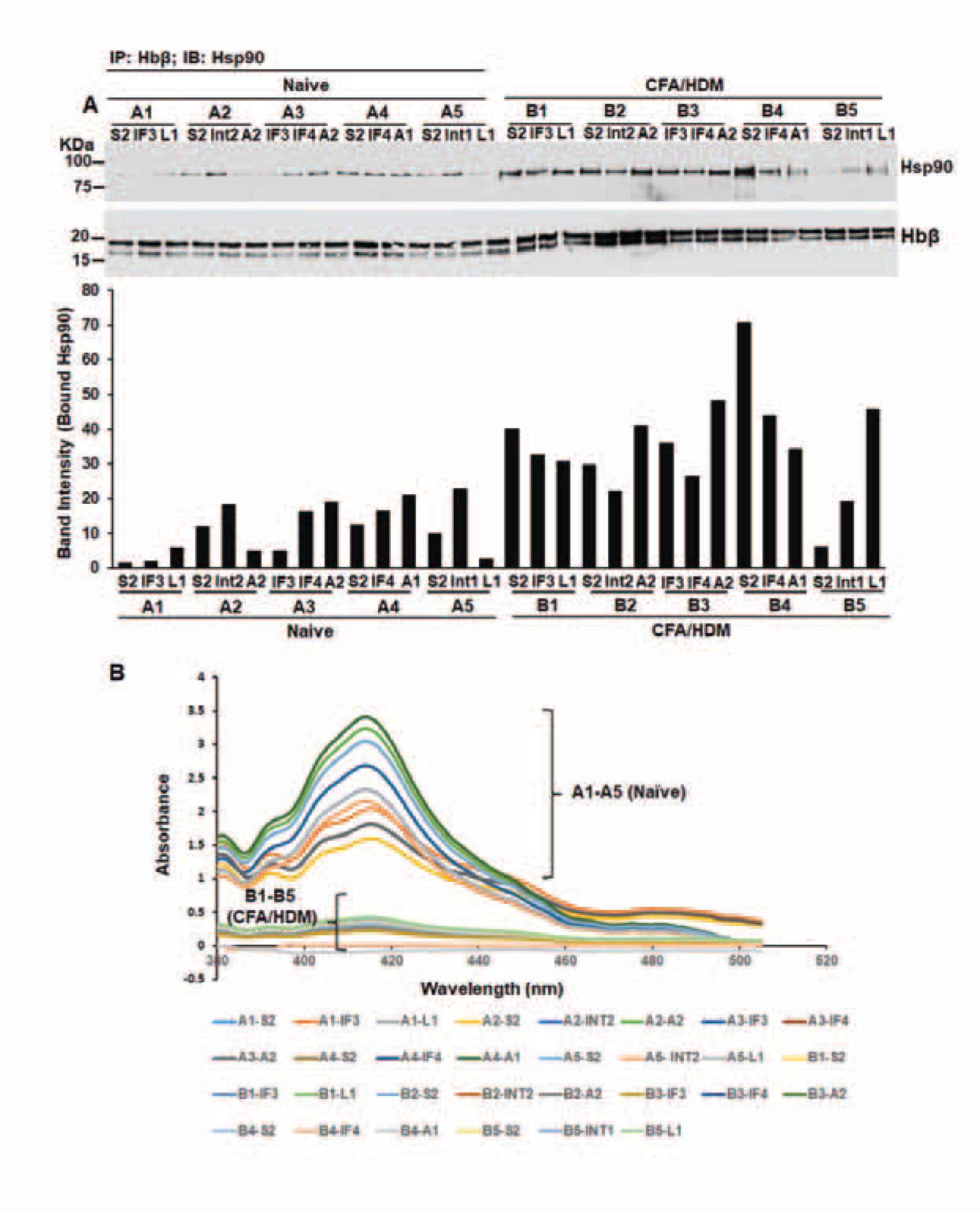
Status of Hb heme in CFA/HDM lung segments. IPs were performed to detect the strength of Hbβ-hsp90 interaction which is a measure of heme-free Hbβ and Hbβ spectra of mouse lung supernatants was estimated by absorption spectra collected with an integrating sphere detector (Olis Clarity), using bead bound Hbβ that was immunoprecipitated with Hbβ antibody. (A) IPs depicting representative Hbβ-hsp90 interactions in various mouse lung segments as indicated and its corresponding densitometries normalized by the bound Hbβ. (B) Absorption spectra of naïve and asthma mouse (CFA/HDM) lung supernatants collected with an integrating sphere detector.

**Figure 9.**
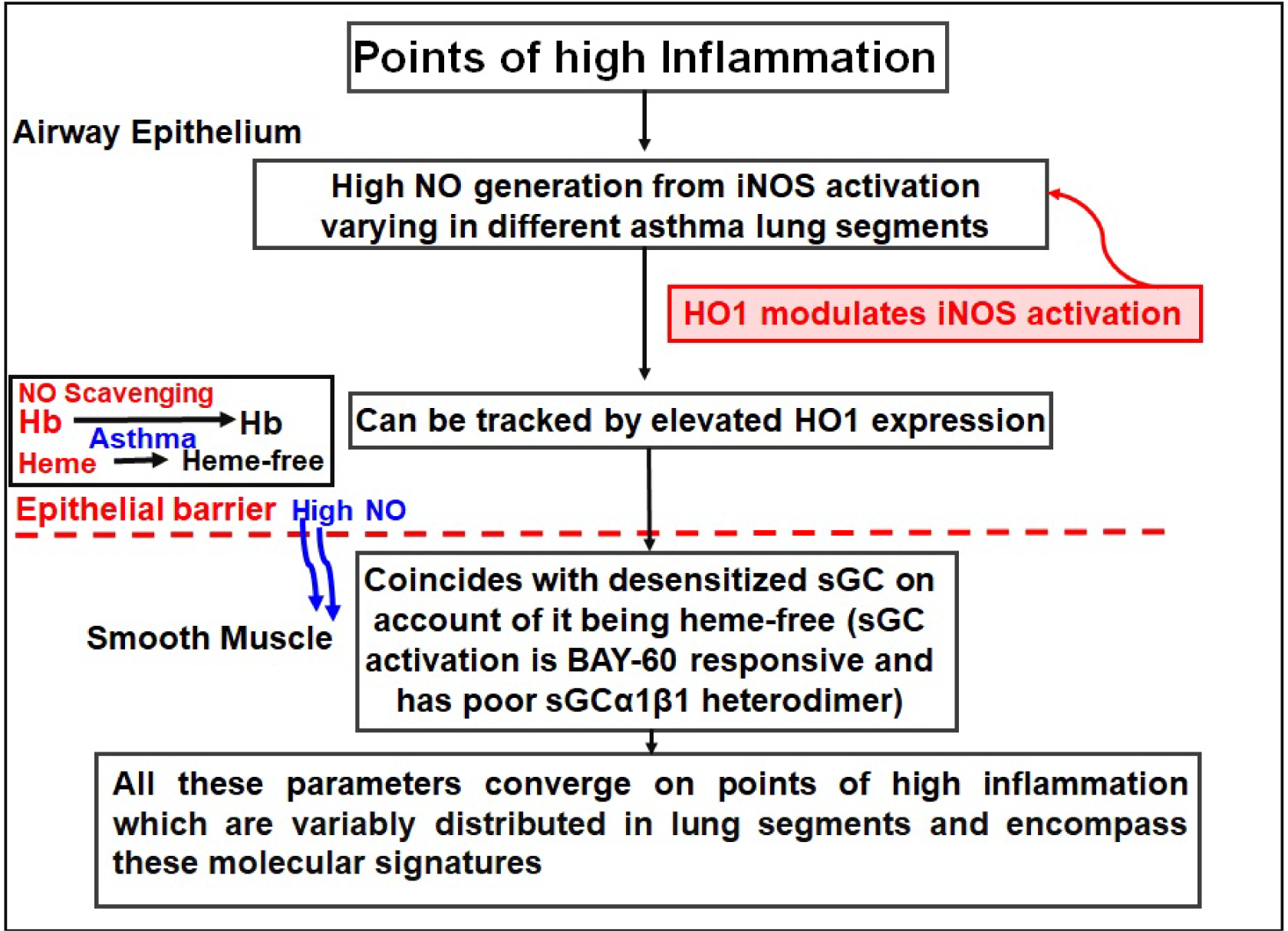
Model establishing the molecular underpinings in mouse asthma between point of high inflammation and sGC dysfunction. Severe inflammation points in mouse asthma (OVA, CFA/HDM) lungs are characterized by high NO generated from iNOS induction and these can be tracked by following upregulated expression of HO1 in the different lung segments. HO1 interacts with iNOS to modulate its function in NO generation. High NO generated in the airway epithelium makes lung hemoglobin (Hb) heme-free which is unable to scavenge the excess NO thereby losing its protection for the smooth muscle sGC. High NO thus makes the smooth muscle sGC heme-free and such regions in the lungs overlap with points of upregulated iNOS and HO1. Various lung segments are characterized by different degrees of inflammation which encompass all these molecular signatures.

## Discussion

Our current studies supports a model where severe inflammation points in mouse asthma lungs are characterized by high NO generation from iNOS induction and these can be tracked by upregulated expression of HO1 in the different lung segments (Fig. 9). This study is the first of its kind to demonstrate the distribution and impact of lung inflammation in asthma to molecular signatures of sGC dysfunction using mouse asthma models (OVA and CFA/HDM), where various lung segments are characterized by different degrees of inflammation that encompass all these molecular signatures. Our study demonstrates that lung segments S1/S2, IF2, L1, L3 and A1 in the OVA and S2, IF3, IF4 and A1/A2 in the CFA/HDM lungs have the greatest impact of inflammation on the molecular signatures of sGC dysfunction (Figs. 3 and 5) (20). The mechanistic underpinnings suggests that high NO generated in the airway epithelium makes lung hemoglobin (Hb) heme-free which is unable to scavenge the excess NO thereby losing its protection for the underlying smooth muscle sGC (Figs. 8 and 9) (19). Extending the role of lung Hb in line with our current studies we envisioned that the Hb heme maybe differently impacted in the various inflammed lung lobes/segments in asthma and the loss of Hb heme may correlate with the generation of heme-free sGC. We therefore investigated the heme status of lung Hb in the CFA/HDM lung segments which gave a poor sGC heterodimer and displayed an elevated HO1 expression. Olis spectral measurements revealed that this Hb was heme-free relative to Hb from naïve segments (Fig. 8), thus authenticating our earlier findings (19). High NO thus makes the epithelial Hb and the smooth muscle sGC, heme-free and such regions overlap with points of upregulated iNOS or HO1. Our model conclusively demonstrates that two different asthma models show a non-uniform distribution of inflammation which suggests that specific localized regions of inflammation exists in asthma. Further the results obtained from our current studies on mouse asthma lungs may further translate into human lungs. However since there are basic structural differences between human and mouse lungs (eg. humans have two lung lobes on the left and three on the right while mouse has one on the left and four on the right) (44) we envision that regions of lung inflammation in the various segments of the human lungs may differ, but the correlations between the parameters of inflammation in terms of iNOS upregulation and subsequent NO production should impact the human smooth muscle sGC in a similar manner as demonstrated by our current study.

HO1 is not only a heme-catabolizing enzyme but also has a protective immunoregulatory role in asthmatic airway inflammation (45). In its protective role many studies have shown that HO1 expression is upregulated in asthma patients (46–48) and in animal models of asthma (49–52). Antioxidant function of HO1 plays a vital role in the maintenance of sGC heme in its reduced state (ferrous), which is responsive to NO as arteries of HO1^-/-^ mice display the heme-free sGC phenotype (53) which is BAY-60 responsive and is similar to the sGC dysfunction caused during allergic asthma (Figs. 3 and 5) (20). HO1 has vital regulatory effects on multiple types of immune cells which participate in the formation of chronic airway inflammation during asthma onset (49, 54–57) and published evidence thus far suggests that HO1 function in asthma is to avoid further worsening of inflammation as inhibition of endogenous HO1 further aggravates inflammation (58, 59). Moreover upregulation of HO1 resulting in CO and bilirubin generation, which are degradation products of heme-catabolism also have significant protective effects on allergic airway inflammation. These factors are known to inhibit plasma exudation to the trachea, bronchi or segmental bronchi, reduce infiltration of inflammatory cells eg. eosinophils, neutrophils, lymphocytes or macrophages around the airways and the BAL (bronchoalveolar lavage), alleviate airway reactivity or mucus secretion (55–57) and decrease the proportion of antigen-specific Th2 (49) or Th17 cells (54, 60) in mediastinal lymph nodes or the spleen to further inhibit allergic airway inflammation. Our observing specific points of HO1 upregulation in mouse asthma lungs (OVA and CFA/HDM) (Figs. 2, 3 and 5), may in-turn indicate enhanced inflammatory regions where the disease is more severe and HO1 upregulation maybe a protective response to the disease.

Our current study presents the mechanisms causing generation of heme-free sGC, the relative proportions of which vary in the different lung lobes. NO generated from iNOS induction (causing lung inflammation) in the airway epithelium is the primary reason for sGC becoming heme-free in the smooth muscle (20) and the extent of this iNOS activity varies in the different lung segments (Fig. 6). Since HO1 is induced in the lung lobes at points of high inflammation we tested the ability of iNOS and HO1 to interact in OVA lungs or in cytokine activated bronchial epithelial cells in immunoprecipitation assays. iNOS linearly interacts with HO1 (Fig. 7A), and HO1 probably modulates iNOS function to cause more NO generation as iNOS bound HO1 in lung lobes of mouse asthma inversely correlates to the formation of a sGC heterodimer (Fig. 7D). Earlier studies indicated that NO induces HO1 expression in mesenglial cells (61) and that both iNOS and HO1 are co-expressed in the glomerular basement membrane where a possible interaction was envisioned (42). Our current study is probably the first example where iNOS has been found to stably interact with HO1 in mouse asthma lung lobes, where the strength of iNOS-HO1 interactions inversely correlates to the sGC heterodimer (Fig. 7) which may suggest that HO1 modulates iNOS activation during inflammation in asthma (Fig. S4). The fact that HO1 expression correlates linearly to iNOS activation (Fig. S4) and strength of iNOS-HO1 interaction peaks with the surge in NO synthesis (Fig. 7A, B) suggests that a strong iNOS-HO1 interaction maybe needed for maximum iNOS activation and subsequent NO production. This function of HO1 realized from our current study is different from its regular function of heme catabolism and it thrusts HO1 in a new light where it may modulate iNOS activity for NO production.

While there is overlap among the inflammatory lung segments from both the OVA and the CFA/HDM models the significance of these studies lies in mapping the molecular underpinings of inflammation. Lung inflammation in the bronchial tissue may drive ventilation heterogeneity in mouse asthma (7) which manifests into infiltration of inflammatory cells eg. eosinophils, neutrophils etc. into the BAL, HO1 upregulation (45), NO generation on account of iNOS generation (20) cascading into generation of NO unresponsive heme-free sGC (20). While neutrophils are sometimes more than eosinophils in the CFA/HDM model, recent studies (62) indicate that it is the eosinophils that drive ventilation heterogeneity in this steroid resistant murine asthma model. This may relate to similarities in the molecular signatures of inflammation in the OVA or the CFA/HDM model that we see in the current study since eosinophilic inflammation also characterizes the OVA model (20, 22, 63). While early or late asthmatic responses (EAR or LAR) can change the location of airway obstruction from central to small airways, it is the more localized regional airflow obstruction caused due to inflammation which mostly impacts the small airways and can extend with the disease progression (2, 5). Further looking at the points of inflammation as it transitions from central to the smaller airways one can use the molecular fingerprints of inflammation combined with the cellular infiltration to predict the type of asthma or the extent of disease severity.

### Future directions

Further studies can aim to evaluate molecular signatures of dysfunction or refractoriness which generates in other cyclase pathway working via β-adrenergic receptors (64) and correlate those parameters with the distribution of lung inflammation. Studies can also aim to generate an algorithm using AI to list the distribution of specific molecular fingerprints of inflammation for lung segments in mouse asthma models. Moreover establishing a similar analysis of correlating molecular markers of inflammation in mouse asthma models with lung ventilation heterogeneity data obtained from 4-Dimensional lung imaging scans (7) will lead to an enhanced understanding of the disease pathogenesis of asthma. Our current study provides a corridor of correlating ventilation heterogeneity with molecular markers of inflammation and these studies can be now be further pursued.

## Abbreviations

AI, Artificial intelligence; A1/A2, Accessory lobes 1/2; IF1-IF4, Inferior lobes (1 to 4); Int1/Int2, Intermediate lobes 1/2; L1-L4, Left lobes (1 to 4); S1/S2, Superior lobes 1/2; BAY-60, BAY 60-2770; BAY-41, BAY 41-2272; BAL, Bronchoalveolar lavage; DAB, Diaminobenzidine; EAR, Early asthmatic responses; LAR, Late asthmatic responses; Mch, Methacholine; OVA, Ovalbumin; CFA/HDM, Complete freund’s adjuvant/House dust mite; FFPE, Formalin-fixed paraffin embedded.

## Data Availability Statement Included in the article

The data that support the findings of this study are available in the methods, results and/or supplementary material of this article.

## Acknowledgements

We acknowledge Alisha Slomers for her assistance with running western blots. This work was supported by National Institute of Health Grants R56HL139564 and R01HL150049 (A.G.)

## Conflict of Interest

The authors declare no conflict of interest.

## Author Contributions

A. Ghosh, R. Westcott, E. Stuehr and C. Ghosh designed the experiments. A. Ghosh, M. P. Sumi, R. Westcott, E. Stuehr and C. Ghosh performed all cell culture and biochemical studies. A. Ghosh, M. P. Sumi, R. Westcott, E. Stuehr, C. Ghosh and D. J. Stuehr analyzed all the data. A. Ghosh wrote the manuscript.

## Supplementary Information

**Figure S1.**
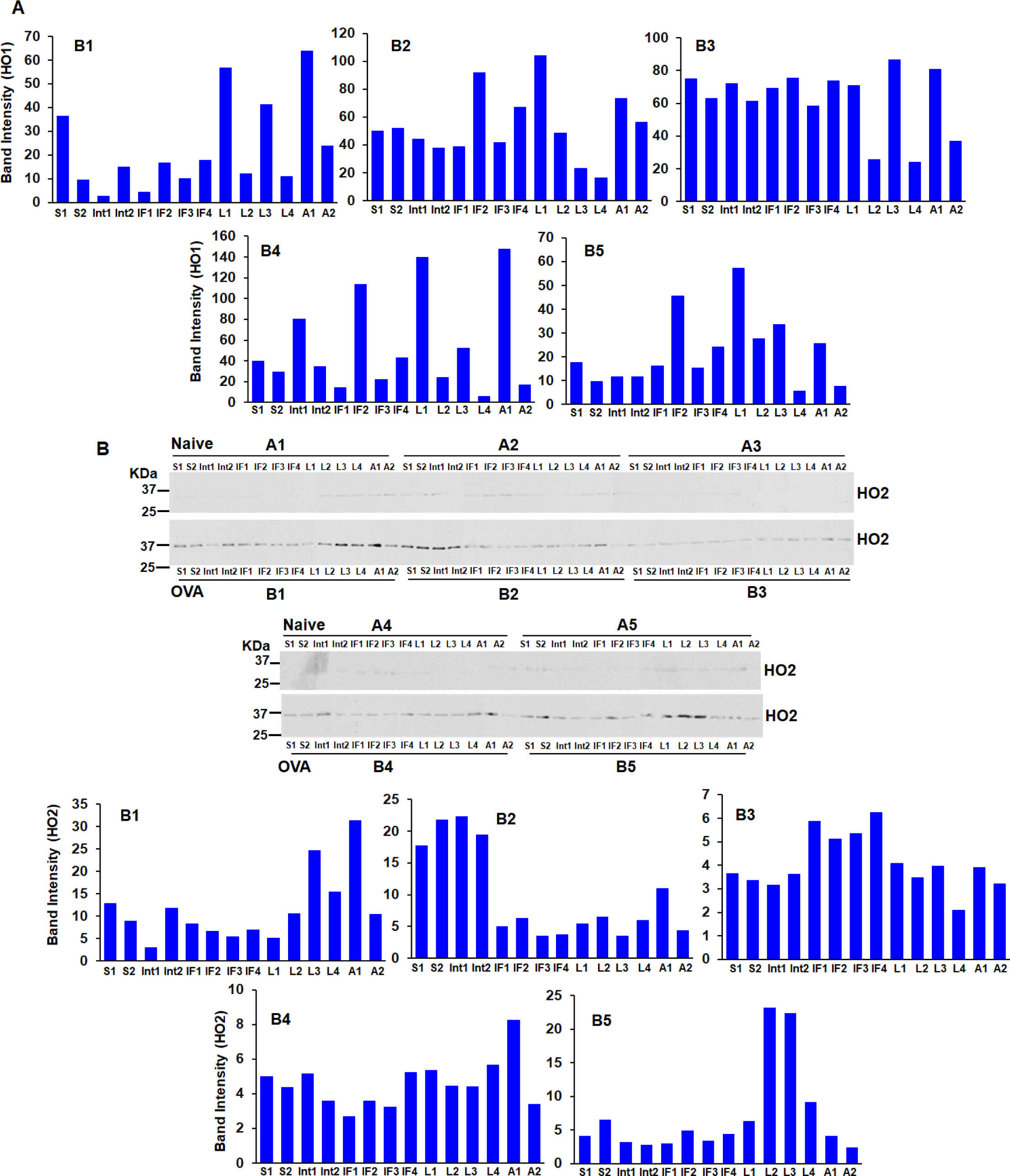
Expression levels of HO2 and densitometries of HO1/HO2 from lung segments of OVA mice (n=5, OVA mice). Mouse lung supernatants generated from all the 14 segments of a mouse lung were run on SDS-PAGE and western blotted with specific antibodies as mentioned in figure 2. (A) Corresponding densitometries of HO1 expression normalized to loading control β-actin as depicted in figure 2. (B) Representative expression HO2 (upper panel) in naïve (A1-A5) and OVA (B1-B5) mice lung segments with its corresponding densitometries (lower panel).

**Figure S2.**
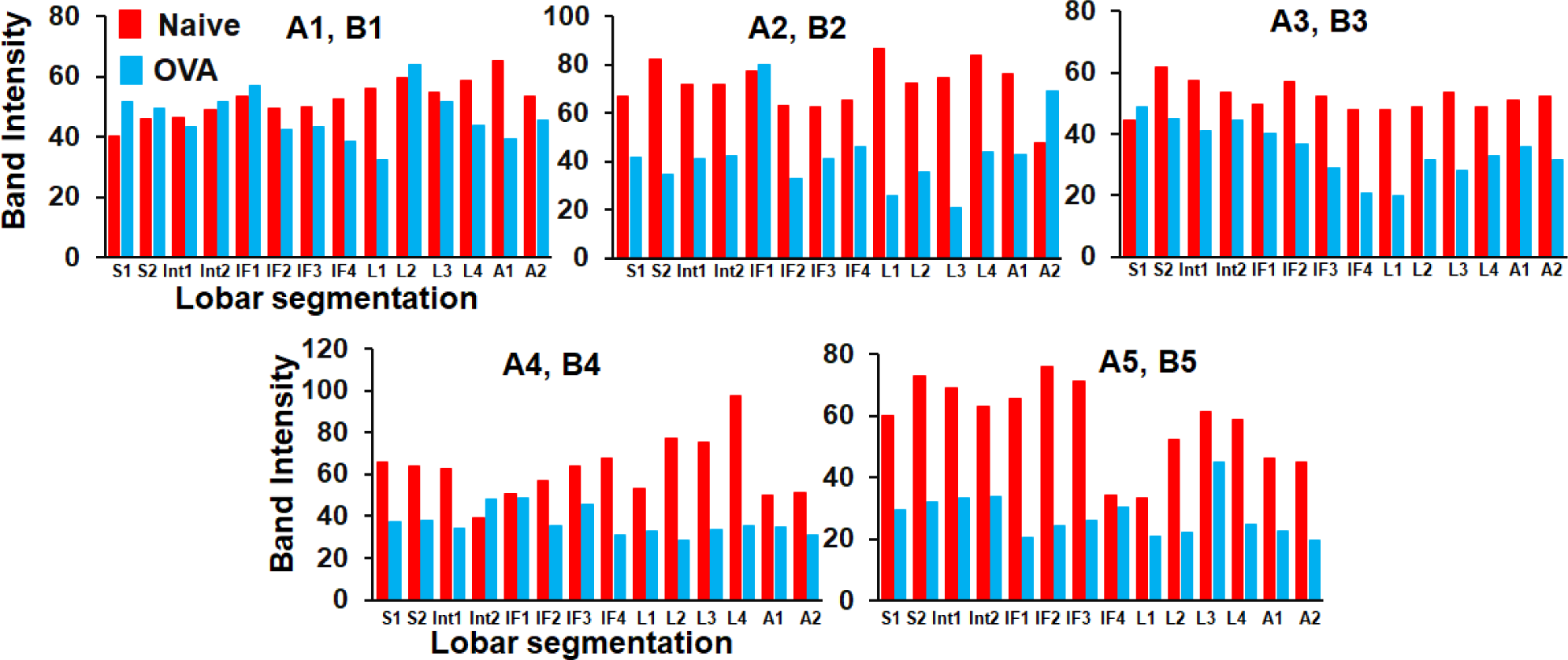
Densitometries of sGCβ1 expression of control naïve and asthma (OVA) mice lung segments derived from corresponding westerns depicted in figure 3A.

**Figure S3.**
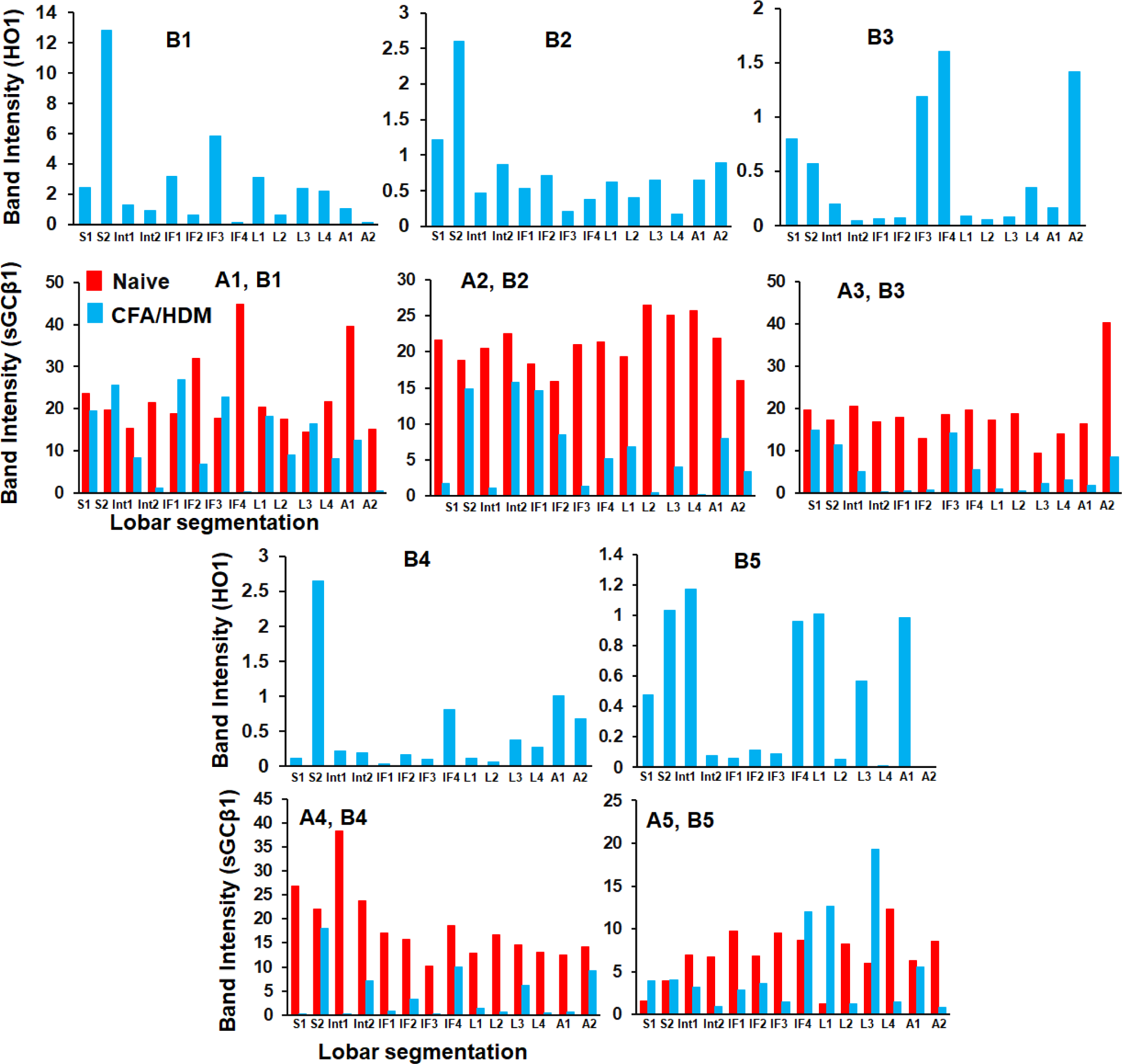
Densitometries of HO1 and sGCβ1 expression in the lung segments from control naïve and asthma (CFA/HDM) mice derived from corresponding westerns depicted in figure 4A.

**Figure S4.**
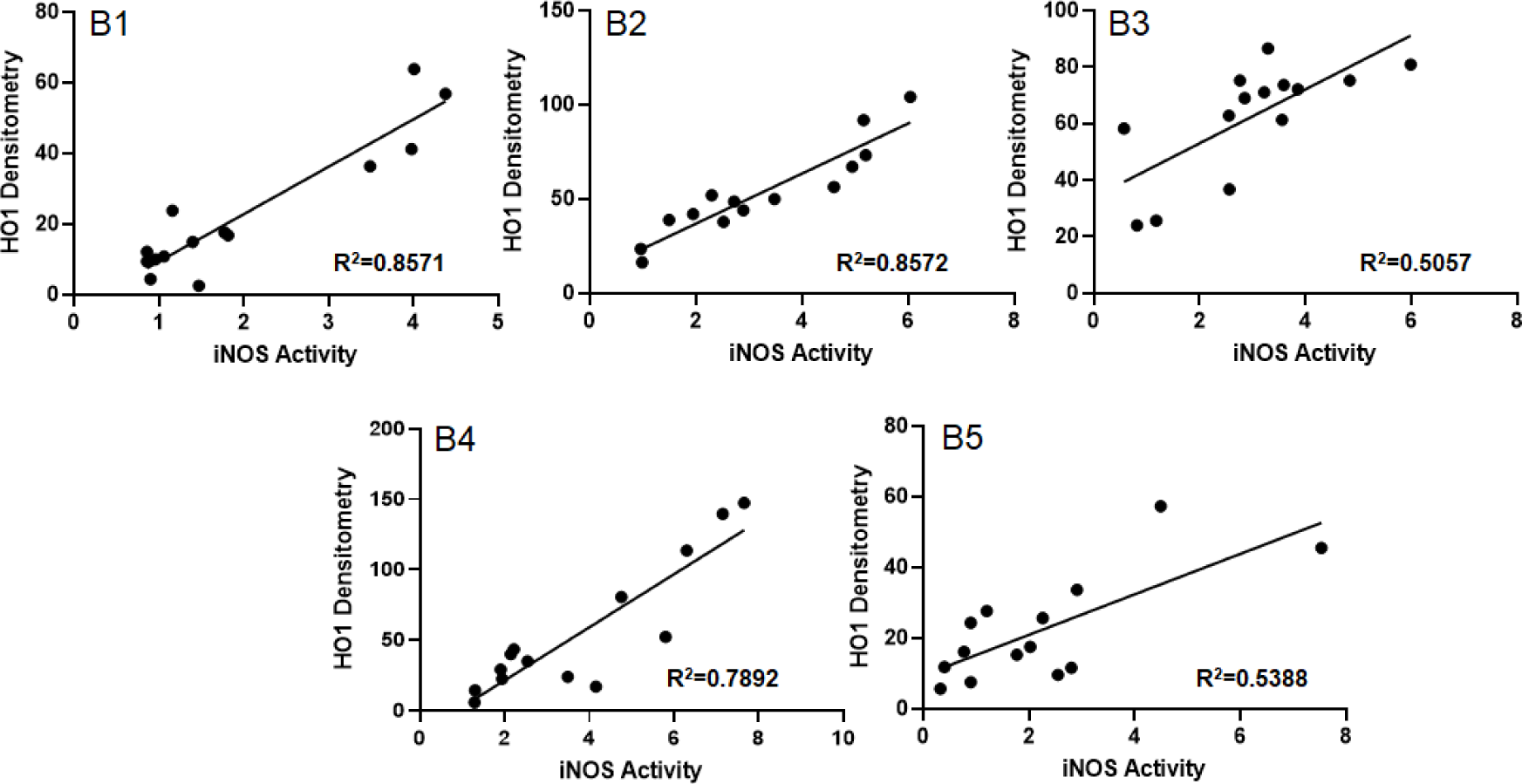
HO1 densitometries as depicted in figure S1 correlated with iNOS activity determined in the lung segments of OVA mice from figure 6 (n=5, OVA mice).

**Figure S5.**
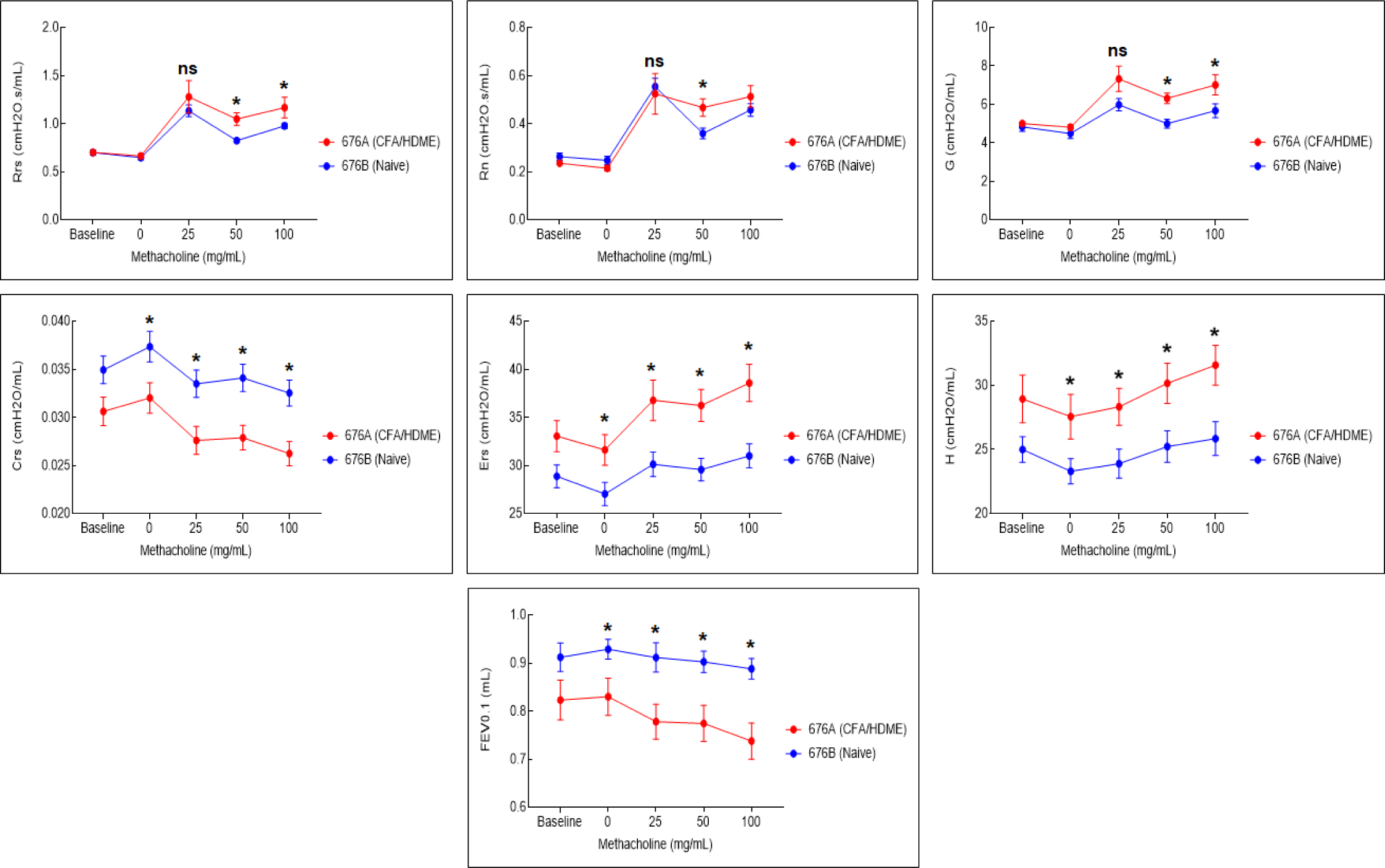
Lung Airway hyper reactivity (AHR) measurements in CFA/HDM asthma mice relative to naïve mice. Airway resistance (Rrs and Rn) and parameters of lung mechanics (G, tissue damping [resistance]; Ers, elastance; Crs, variables compliance; H, tissue elasticity) were measured on a flexivent in response to increasing doses of methacholine (Mch) used as a bronchoconstrictor in CFA/HDM asthma and naïve mice (n=6 each for asthma or naive). Data are mean (n= 6, CFA/HDM mice) ± SD. *p < 0.05, by one-way ANOVA. ns is statistically non-significant.

**Figure S6.**
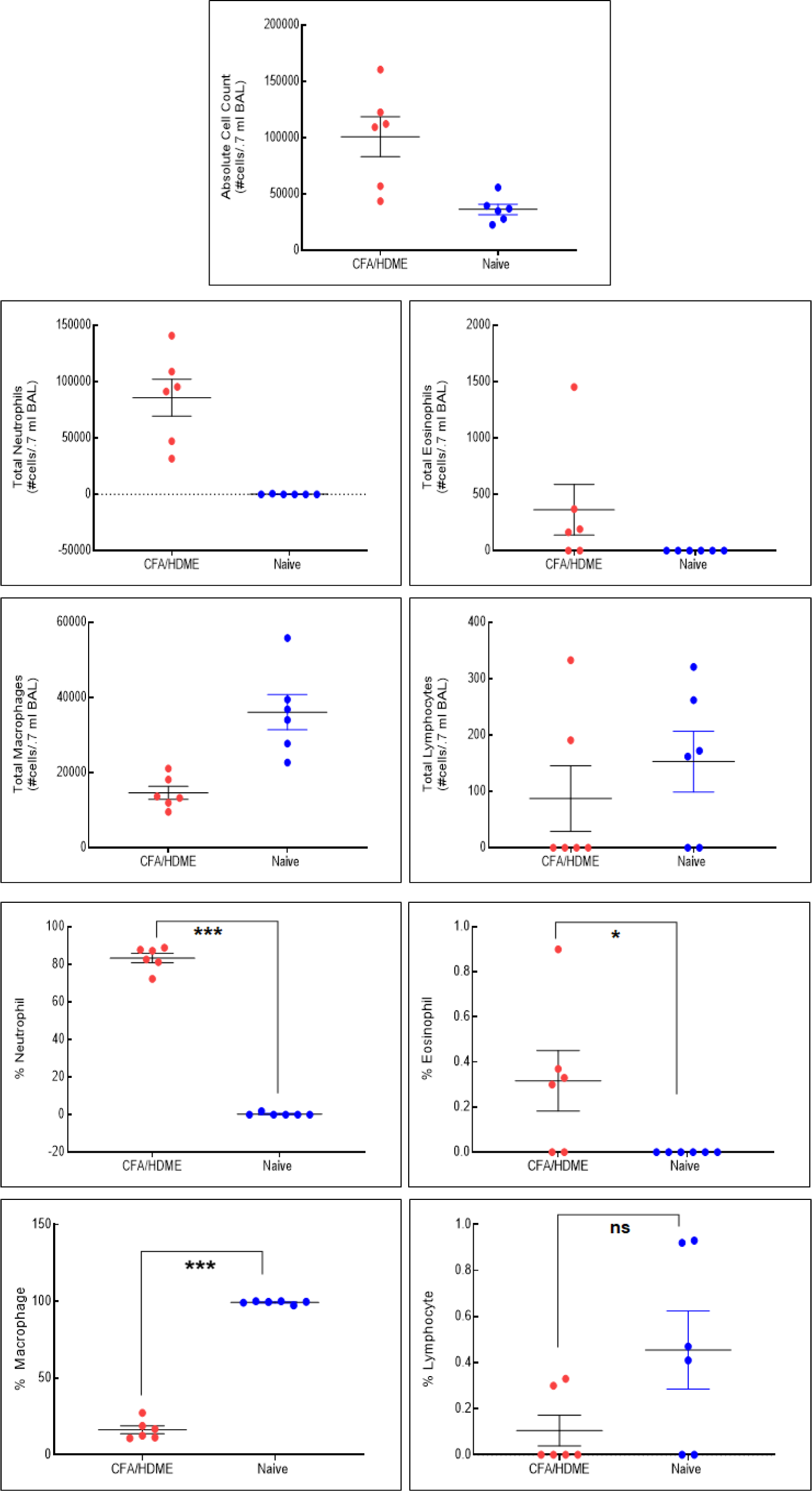
Changes in mouse lung associated with inflammatory asthma. Total cell counts and calculated percentages of neutrophils, eosinophils, macrophages and lymphocytes in bronchial lavage fluid from naïve and from mice with allergic asthma toward CFA/HDM (n = 6). Data are mean (n= 6, CFA/HDM mice) ± SD. *p < 0.05, **p < 0.01, ***p < 0.001 by one-way ANOVA. ns is statistically non-significant.

**Figure S7.**
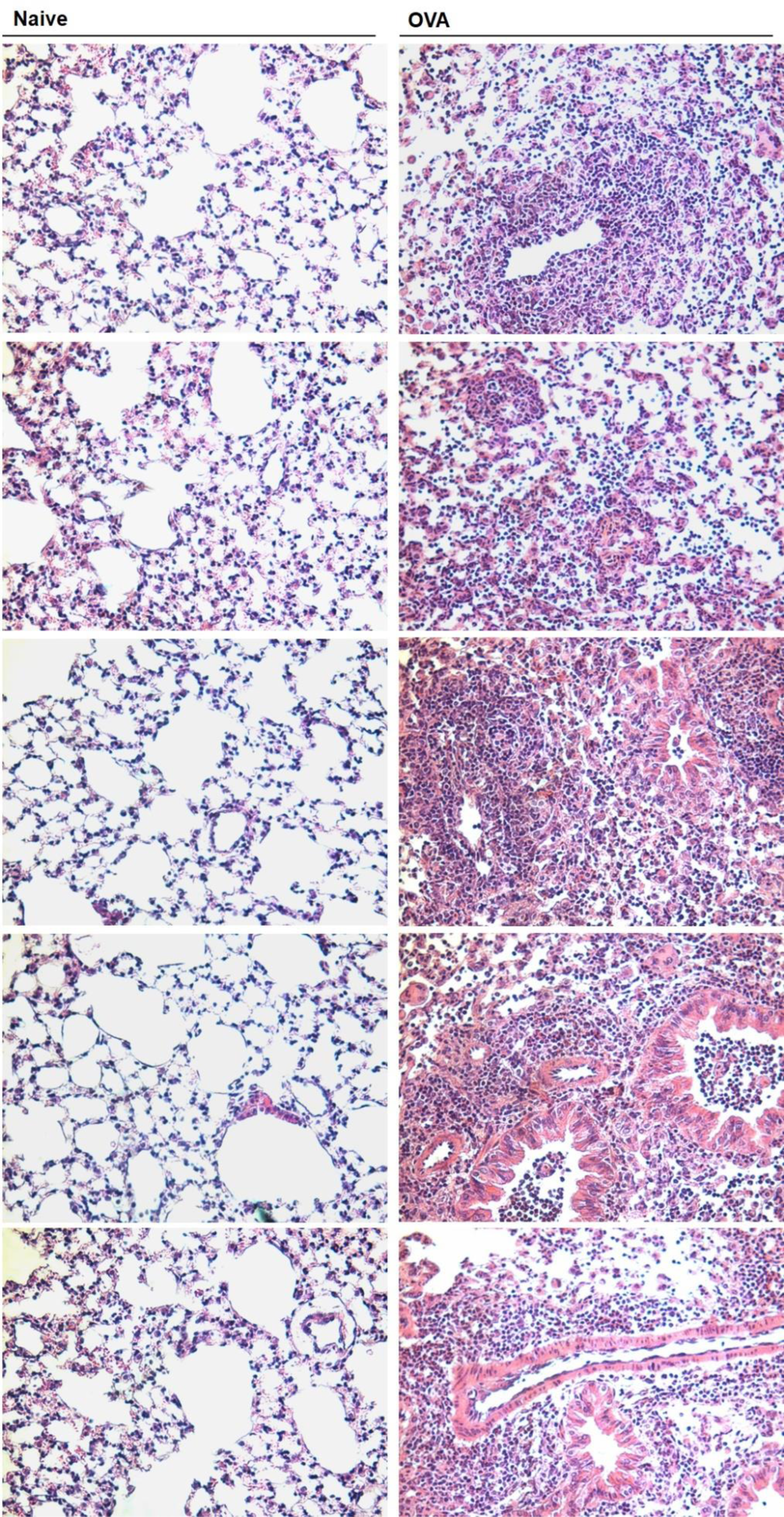
H & E staining of naïve and OVA lung sections depicting tissue histology, captured in bright field at 20X. Representative images from Naïve (left) and OVA (right) lung sections arranged lengthwise. OVA lung sections depict reduction in lung airways from inflammation and infiltration of cells.

**Supplementary Table S1.** Antibodies used and their sources.

| **Serial No.** | **Antibody** | **Species-Clonality** | **Source (Catalog No.)** |
| --- | --- | --- | --- |
| 1. | iNOS | Mouse monoclonal | Invitrogen (VF3008463) |
| 2. | iNOS | Rabbit polyclonal | Cell Signaling Tech. (39898) |
| 3. | iNOS | Rabbit polyclonal | Abcam (ab15323) |
| 4. | sGCβ1 | Rabbit polyclonal | SIGMA, ER-19 (G4405) |
| 5. | sGCβ1 | Rabbit polyclonal | Cayman Chem.(160897) |
| 6. | sGCα1 | Rabbit polyclonal | Abcam (ab101368) |
| 7. | Trx1 | Rabbit polyclonal | Origene (TA322865) |
| 8. | Smooth muscle actin | Mouse monoclonal | Santa Cruz (sc-53015) |
| 9. | Smooth muscle actin | Rabbit polyclonal | Cell Signaling Tech.(D4K9N) |
| 10. | β-Actin | Mouse monoclonal | SIGMA (A5441) |
| 11. | HO1 | Rabbit polyclonal | Cell Signaling Tech. (5061S) |
| 12. | HO1 | Mouse monoclonal | Invitrogen (WF3315691) |
| 13. | HO1 | Rabbit polyclonal | Cell Signaling Tech. (70081) |
| 14. | HO2 | Rabbit polyclonal | Cell Signaling Tech. (32790S) |
| 15. | Catalase | Mouse monoclonal | Santa Cruz (sc-271803) |

## Notes

### Competing Interest Statement

The authors have declared no competing interest.

